# Id proteins suppress E2A-driven innate-like T cell development prior to TCR selection

**DOI:** 10.1101/208728

**Authors:** Sumedha Roy, Amanda J. Moore, Cassandra Love, Anupama Reddy, Deepthi Rajagopalan, Sandeep S. Dave, Leping Li, Cornelis Murre, Yuan Zhuang

## Abstract

Id proteins have been shown to promote the differentiation of conventional αβ and γδT cells, and to suppress the expansion of invariant Natural Killer T (iNKT) cells and innate-like γδNKT within their respective cell lineages. However, it remains to be determined whether Id proteins regulate lineage specification in developing T cells that give rise to these distinct cell fates. Here we report that in the absence of Id2 and Id3 proteins, E2A prematurely activates genes critical for the iNKT cell lineage prior to TCR expression. Enhanced iNKT development in Id3-deficient mice lacking γδ NKT cells suggests that Id3 regulates the lineage competition between these populations. RNA-Seq analysis establishes E2A as the transcriptional regulator of both iNKT and γδNKT development. In the absence of pre-TCR signaling, Id2/Id3 deletion gives rise to a large population of iNKT cells and a unique innate-like DP population, despite the block in conventional αβ T cell development. The transcriptional profile of these unique DP cells reflects enrichment of innate-like signature genes, including PLZF (Zbtb16) and Granzyme A (Gzma). Results from these genetic models and genome-wide analyses suggest that Id proteins suppress E2A-driven innate-like T cell programs prior to TCR selection to enforce predominance of conventional T cells.

## Introduction

The thymic output of a diverse and abundant population of conventional CD4^+^ and CD8^+^ αβ T cells constitutes the adaptive immune system that is necessary for a specific and effective immune response to antigens. A smaller but significant population of unconventional T cells also concomitantly develops in the thymus, with innate-like capabilities of mounting a non-specific but rapid immune response to antigens. These innate-like T cells have garnered increasing interest in lieu of their memory phenotype that can be harnessed in the context of allergies, infections and tumors. They often possess limited TCR diversity that allows recognition of self ligands presented by non-classical MHC molecules such as CD1 and MR1.^[1]^ They are also characterized by high levels of expression of the innate-like transcription factor, Promyelocytic Zinc Finger (PLZF).^[2–6]^ While the transcriptional programs that drive conventional CD4^+^ and CD8^+^ T cell specification and development have been well characterized, little is known about the innate-specific transcriptional programs upstream of PLZF that are responsible for the divergence of innate-like T cells from conventional T cells.

Innate-like T cell populations include TCRαβ^+^ Natural Killer T (NKT) cells, TCRγδ^+^ NKT cells, innate-like CD8^+^ T cells, CD8αα intraepithelial lymphocytes (IELs), and Mucosal-associated invariant T (MAIT) cells. Invariant NKT (iNKT) cells are among the best characterized innate-like T cells, which arise in parallel with conventional αβ T cells. These cells are known to stochastically express a canonical Vα14-Jα18 TCRα chain at the CD4^+^CD8^+^ double positive (DP) stage, which allows them to undergo TCR selection mediated by a CD1d molecule expressed on other conventional DP thymocytes.^[7, 8]^ γδNKT cells are yet another population of innate-like T cells that express a restricted Vγ1.1Vδ6.3 TCR. Both iNKT and γδNKT cells express high levels of PLZF and readily produce effector cytokines like IL-4^[2, 9]^.

Id proteins, primarily produced by *Id2* and *Id3* during T cell development, are inhibitors of the E protein transcription factors E2A and HEB. Interestingly, Id proteins play opposite roles in the development of conventional and innate-like T cells, such that they promote the former and suppress the latter. In response to pre-TCR and TCR signals, Id proteins inhibit E protein activity to play a critical role in promoting the differentiation and positive selection of conventional αβ T cells.^[10–12]^ The Id-mediated inhibition is both necessary and sufficient to activate gene expression programs critical for T cell maturation. In line with this, it has been shown that disruption of *Id2* and *Id3* impairs conventional αβ T cell development beyond the TCR checkpoint^[13]^. Analogous to αβ T cell development, the function of Id3 in promoting conventional γδ T cell development has also been mapped downstream of the γδ TCR^[14]^. In contrast, however, large populations of iNKT, γδNKT and innate variant T_FH_ cells have been observed in the same *Id3*- and *Id2/Id3*-deficient animals, indicating a negative role for Id proteins in regulating innate-like T cell development.^[15–21]^ However, the mechanism that drives the development and expansion of these innate-like T cell populations in *Id*-deficient mice is still elusive. Given the contradictory nature of Id proteins in supporting conventional T cells and suppressing innate-like T cells, it is reasonable to predict that Id proteins control innate-like T cell development through a distinct, or additional mechanism, that is TCR signaling-independent. Id proteins have also been shown to modulate E protein activity throughout T cell development.^[10]^ Therefore, it remains to be determined whether Id-mediated suppression of these innate-like T cells is limited to their expansion upon selection and lineage commitment, or if it also influences their lineage specification prior to TCR selection.

We first investigated the role of E proteins, particularly E2A, in iNKT cell development in the absence of Id proteins. E2A ChIP-Seq revealed several genes that are important for iNKT cell development, such as *Tcff*^*[22]*^, *Egr2*^*[23,24]*^ and *Bcl11b*^*[25]*^, to be direct targets of E2A. This suggested that Id proteins suppress E2A-driven iNKT-specific genetic programs. We then examined any developmental competition between iNKT and γδNKT cells in *Id3*-deficient mice, which have enhanced development of both these lineages.^[20, 21]^ In the absence of γδ T cells, *Id3*-deficient mice had increased iNKT commitment and development, supporting a role for Id3 in regulating the outcome of lineage competition between γδNKT and iNKT cells. Systematic analysis of the transcriptional profiles of expanded iNKT and γδNKT populations, along with the genome-wide E2A binding data, demonstrated that E2A drives downstream transcriptional programs that support both iNKT and γδNKT development in the absence of Id protein(s). We further explored the impact of *Id* deficiency on innate-like T cell development at an earlier stage, i. e. prior to the pre-TCR checkpoint. We found that the deletion of Id proteins in pre-Tα^−/−^ mice promoted iNKT cell development without rescuing conventional αβ lineage development. Furthermore, this lack of Id proteins in the absence of pre-TCR signaling uniformly turned on an innate-like transcriptional program among all T cells, including DP and γδ T cells. Collectively, these studies provide compelling evidence to demonstrate the role of Id proteins in the suppression of iNKT and innate-like developmental programs through inhibition of E protein activity, prior to both TCR and pre-TCR selection checkpoints. Our findings also suggest that this suppression mechanism is essential for the predominance of conventional T cell development in the thymus.

## Materials and Methods

### Study design

Appropriate sample size was determined empirically for each individual experiment or technique in a manner such that statistical significance is reached and scientific rigor is achieved with minimum number of mice necessary. For most experiments, 3 to 5 mice were sufficient per genotype to statistically observe dramatic phenotype changes. Certain experiments depicting cell numbers are cumulative, and have greater number of mice. Each figure is representative of at least 3 sets of independent experiments. All references of replicates in this study are indicative of biological replicates, i.e. individual animals, and not technical replicates. All data points obtained are included in each experiment without any exclusion.

### Mice

Id2^f/f^Id3^f/f^LckCre^+^ (L-DKO), Id3^−/−^ and Id3^−/−^ TCRδ^−/−^ mice were generated as previously described.^[15, 26]^ L-DKO pTα^−/−^ mice were generated by breeding L-DKO mice with pTα^−/−^ mice,^[27]^ which were a generous gift from David L. Weist (Fox Chase Cancer Center, Philadelphia). All mice were bred in a specific pathogen-free facility of Duke University Division of Laboratory Animal Resources, and all procedures were performed according to protocols approved by the Institutional Animal Care and Use Committee.

### Cell sorting and RNA extraction

All cells were sorted in FACS buffer using a MoFlo XDP cell sorter. Total mRNA from sorted cells was extracted using an RNAqueous Kit (Life Technology) according to manufacturer's protocol.

### Flow cytometry

Surface marker antibodies were used according to manufacturer's protocol (Biolegend). Intracellular staining with PLZF antibody (eBioscience) was done using the Foxp3 staining buffer kit (eBioscience). CD1d tetramers were received from the Tetramer Facility of the National Institutes of Health. Stained samples were run on a FACSCanto II machine (BD Biosciences) and data was further analyzed with FlowJo software (Tree Star). Bar graphs were drawn using GraphPad Prism (GraphPad Software). Two-tailed student’s t-test was used for statistics, with p values less than 0.05 considered significant.

### ChIP-Seq analysis

26 x 10^6^ iNKT and 30 × 10^6^ DP cells were sorted and pooled from multiple L-DKO mice for the E2A ChiP-Seq analysis. 2 × 10^6^ γδ T cells were also sorted from Id3^−/−^ mice for H3K4me2 analysis. iNKT (CD1dTet^+^ TCRβ^+^) and DP (CD1dTet^−^ CD4^+^ CD8^+^) cells were sorted from 3-5 weeks old L-DKO mice. For H3K4me2 ChIP-Seq, cells were fixed with 1% formaldehyde, whereas for E2A ChIP-Seq, cells were additionally fixed with 1.5 mM EGS (ethylene glycol-bis(succinic acid N-hydroxysuccinimide ester)). Crosslinked cells were lysed, nuclei were extracted, and sonicated using Bioruptor Plus (Diagenode) and immunoprecipitated with E2A (V-18, Santa Cruz Biotechnology, Lot G0814) or H3K4me2 (Millipore, 07-030, Lot 2430486) antibody. After elution and reverse crosslinking, RNA and proteins were digested, followed by DNA purification using a ChIP DNA Clean and Concentrator kit (Zymoresearch). Libraries were prepared with the NEBNext primer set, which included applying ChIP DNA to end repair, A-tailing, adapter ligation and PCR amplification. Samples were cleaned and size selected by 8% PAGE or AMPure beads (Agencourt). Sequencing was done on HiSeq4000 platform (Illumina). ChIP-Seq sequenced reads were aligned to the mm9 genome using Bowtie^[28]^ software (version 1.1.2, parameters: --chunkmbs 128 --mm -m1 --best --strata -p4 -S -q). Peaks were called using MACS^[29]^ (version 1.4.2, default parameters). Peaks were then annotated by the NGS: Peak Annotation tool on Nebula.^[30]^ Bed and wiggle files were generated by MACS for visualization using the Integrative Genomic Viewer.^[31]^ De-novo motif analysis was done using the findmotifs.pl program with HOMER^[32]^ (v4.7.2, 50 or 200 bp within each peak).

### RNA-Seq analysis

iNKT (CD1dTet^+^ TCRβ^+^), γδNKT (TCRγδ^+^ CD3^+^) and DP (TCRγδ^−^ CD4^+^ CD8^+^) cells were sorted from 4-5 weeks old WT (total 2 × 10^5^ γδ T, 2 × 10^5^ iNKT cells, 2 × 10^6^ DP cells), Id3^−/−^ (total 15 × 10^6^ γδ T cells) or L-DKO (total 15 × 10^6^ iNKT cells, 15 × 10^6^ DP cells) mice. After RNA extraction using the RNAqueous kit, and quality was assessed using the Agilent Bioanalyzer RNA Pico chip. Ribosomal RNA was depleted using the RiboErase method from Kapa Biosystems. In short, 1ug of total RNA was hybridized with 1ug of hybridization oligos tiling the 18s, 28s, 5.8s and mitochondrial rRNA sequences. Each sample was then RNaseH treated to degrade complementary rRNA sequence. The product was cleaned and purified using 2.2X AMPure beads (Agencourt). The cleaned product was DNase treated to degrade the DNA oligo mix. The remaining rRNA depleted samples were then purified using 2.2X AMPure XP beads. The Kapa Stranded RNA-Seq Kit was used to generate stranded Illumina sequencing libraries (Kapa Biosystems). RNA from was fragmented at 94°C for 6 minutes. Briefly, RNA was hybridized to random primers, followed by first-strand cDNA synthesis, second-strand cDNA synthesis with marking, A-tailing, ligation of Illumina paired-end adapters with 8 bp barcodes, and 9 cycles of PCR amplification. Reactions were purified with Agencourt AMPure XP beads where necessary. Libraries were multiplexed in equimolar amounts, and sequenced as paired-end 50-bp reads using a HiSeq2500 platform (Illumina).

A second, independent round of RNASeq was done with DP (TCRγδ^−^ CD4^+^ CD8^+^) cells sorted from 5 weeks old WT (9 × 10^5^ cells), pTαKO (4.5 × 10^5^ cells), L-DKO (9 × 10^5^ cells) and L-DKO pTαKO (9 × 10^5^ cells) mice. 150 bp paired-end sequencing was done on the HiSeq2500 platform (Illumina). All subsequent data analysis was done using the steps outlined above. RNA-Seq sequencing reads were first trimmed using Trimmomatic.^[33]^ Read alignment was done using Tophat and expression quantification was done using Cufflinks.^[34]^ Log2 transformed FPKM (fragments per kilobase exon-model per million reads mapped) were used for downstream analyses. Further filtering of low quality genes, Principal Component Analysis, statistical analysis and visualizations were done using R.^[35]^ Pathway analysis was done using the Molecular Signatures Database (MSigDB) v5.2^[36]^.

### Innate gene signature and network analysis

Raw microarray expression data was requested and downloaded from Immgen for selected subsets: preT_DN3A_Th (DN3a), preT_DN3B_Th (DN3b), T_DN4_Th (DN4), T_DP_Th (DP), T_4SP69+_Th (post-selection CD4SP), NKT_44-NK1_1-_Th (stage 0 and 1 iNKT cells), Tgd_Th (total thymic γδ T cells), Tgd_vg1+vd6+24ahi_Th (immature Vγ1.1V56.3 cells). Average gene expression among DN3a, DN3b, DN4, DP and CD4SP cells was assumed to be the reference conventional αβ T cell population. Total thymic γδ T cells were considered as reference for conventional γδ T cell population. Fold change in expression for iNKT and γδNKT cells was calculated with respect to the reference conventional αβ and γδ T cell populations respectively. Genes that had more than 1.5 fold upregulation or 0.6 fold downregulation among <u>both</u> iNKT and γδNKT cells were considered to represent the “innate-like gene signature”. These moderately relaxed fold change parameters allowed us to ensure that maximal numbers of appropriate genes were captured in this analysis. 189 genes were therefore identified from these specific expression patterns among WT iNKT and γδNKT cells. Additionally, we also included 7 other genes - Tcf3 (E2A), Id2, Id3, Lef1, Sox13, Blk and Sox4 - which have been reported to play important roles in iNKT and γδNKT lineage development, but did not have expression patterns that fit our criteria, i.e. being significantly upregulated or downregulated in both cell types as compared to reference populations. The total 197 genes constituted our innate-like gene signature, derived from Immgen and literature.

111 of the 197 signature genes were found to be dysregulated in Id3^−/−^ γδ and/or in L-DKO iNKT cells. Other known interactions between these 111 genes were retrieved from GeneMania.^[37]^ 83 of the 111 genes were also identified as E2A targets, which had E2A binding to the enhancer, promoter, intragenic, intergenic or downstream regions of these genes, as annotated by Nebula. These interactions, ChIP-Seq targets and gene expression patterns of the 111 genes were represented as a network using Cytoscape3.4.0^[38]^.

### Gene Set Enrichment Analysis (GSEA)

The GSEA^[36, 39]^ desktop application (v2.0) was used to analyze the log2FPKM expression patterns in L-DKO and L-DKO pTα DP samples. 9245 genes that were unchanged in L-DKO DP samples as compared to WT DP samples were included in this analysis. Enrichment in L-DKO pTα DP samples over L-DKO samples was determined using weighted, log2(ratio of classes) parameters and 1000 permutations. The innate-gene signature included 196 genes derived from Immgen, and listed in Supplementary Table 2. The iNKT development and maturation gene set^[40]^ (Msigdb gene set M18517) and inflammatory responses gene sets (Msigdb gene set M5932) were downloaded from Msigdb and used as is.

### Correlation analysis

To determine correlation with Zbtb16 expression, Pearson and Spearman correlation coefficients were determined for all genes across six samples, including replicates of WT DP, pTαKO DP, L-DKO pTαKO DP and L-DKO DP, as derived from RNA-Seq analysis. Genes with both coefficients greater than or equal to 0.7 were considered to be positively correlated, and those with both coefficients less than or equal to −0.7 were considered to be negatively correlated with Zbtb16 expression. Scatter plots were generated using a custom R script.

## Results

### E2A regulates downstream transcription factors critical for iNKT cell development

Several studies have previously demonstrated that the lack of Id proteins in Id2^f/f^ Id3^f/f^ LckCre^+^ (L-DKO, or LckC re-mediated double knockout) mice gives rise to a significantly large iNKT cell population.^[41]^ Therefore, we hypothesized that uninhibited E2A activity may induce genes important for the iNKT developmental program. We performed E2A ChIP-seq analysis using L-DKO DP and L-DKO iNKT cells as representative populations prior to, and after CD1d-mediated selection respectively. We found that the number of E2A peaks in L-DKO DP cells was much greater than in L-DKO iNKT cells (Figure 1a). However, many of the E2A targets in L-DKO iNKT cells were also occupied by E2A in DP cells. These shared targets likely drive and sustain iNKT fate during and after TCR selection, such that E2A is continually bound in both DP and iNKT cells. Next, we performed de novo motif analysis of E2A binding sites to identify transcription factors that potentially cooperate with E2A in regulating the iNKT developmental program (Figure 1b). These identified factors included RUNX1, TCF7, LEF1 and GATA3, which are known to be critical for iNKT development^[8, 22, 42, 43]^. RORγt, which is important for regulating survival of DP cells and, as consequence, for the distal iNKT TCRα rearrangement,^[44]^ was found to be a co-factor only in DP cells, but not in iNKT cells. ChIP-Seq analysis also revealed E2A peaks in L-DKO DP cells at these cooperating transcription factor loci, indicating that E2A can directly regulate these factors to support the downstream iNKT program (Figure 1c). E2A peaks at various loci, including at RORγt, were also shared between our mutant L-DKO cells and previously characterized peaks in DN3, DN4 cells with Wild Type (WT) E protein activity and a T cell line with E protein overexpression, suggesting that E2A might regulate similar targets in developing T cells in WT mice (**Supplementary Figure 1a**)^[11]^.

**Figure 1.**
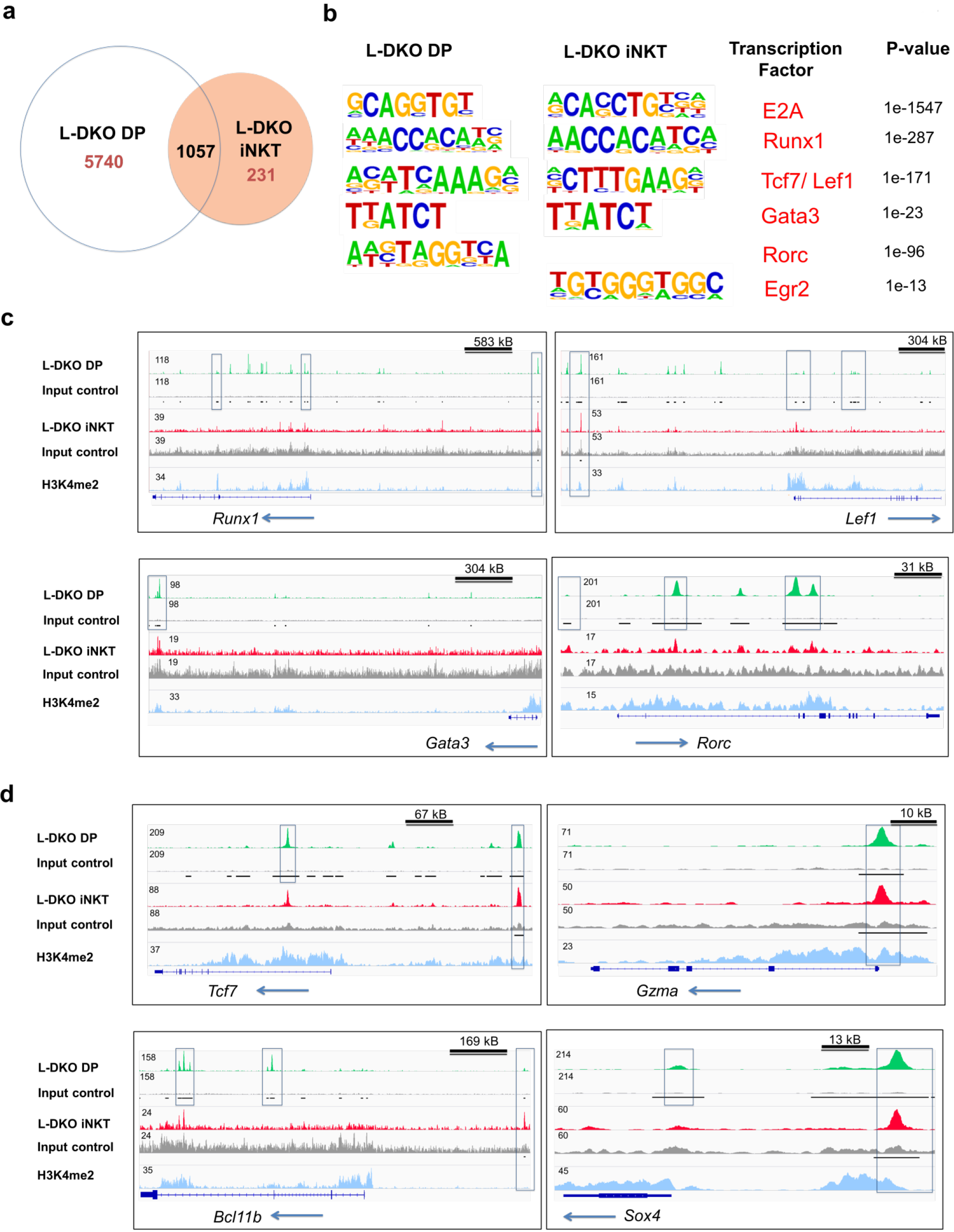
E2A activates and collaborates with transcription factors critical for iNKT cell development. (a) Number of genes with E2A binding in L-DKO DP and/or L-DKO iNKT cells. (b) De novo motif analysis of E2A peaks in L-DKO DP or iNKT samples with p-values (corresponding to DP samples for shared motifs, or for respective samples for unique motifs). (c) E2A peaks in L-DKO DP and L-DKO iNKT cells at loci for motifs identified in (b). (d) E2A peaks at selective loci. Tracks include L-DKO DP and iNKT samples, along with their input control (i.e. not immunoprecipitated with E2A antibody), and H3K4me2 methylation in Id3^−/−^ γδ T cells, which is indicative of transcription factor binding. Important E2A peaks are highlighted by boxes within each panel. Solid black lines below the input control tracks indicate significant (p value less than 10^−5^) peaks called by MACS. Length of the genome depicted in each panel is indicated on the top right. Numbers at the top of each track indicate the maximum peak height.

E2A also bound to the promoter and/or enhancer regions of several genes that have been reported to be critical for iNKT cell development and function (Figure 1d). This included genes such as Tcf7, Bcl11b, Sox4 and Gzma.^[22, 25, 45]^ While direct regulation of PLZF (encoded by Zbtb16) by E2A has been reported previously,^[46]^ we found E2A binding to intragenic, rather than putative regulatory regions of Zbtb16 in L-DKO iNKT or DP cells (**Supplementary Figure 1b**). It is possible that there is an indirect regulation of PLZF through another downstream E2A target in this context. We found E2A bound to the Egr2 promoter (**Supplementary Figure 1b**), which has also been shown to drive PLZF expression in iNKT cells, making it is a likely mediator.^[24]^ Overall, these findings indicate directed regulation of the iNKT developmental program by E2A and other cooperating transcription factors in the absence of Id proteins. More importantly, the comparison of E2A targets between L-DKO DP and L-DKO iNKT cells suggest a possible engagement of E2A in orchestrating an iNKT transcription signature prior to TCR expression.

### Absence of γδ T cells enhances iNKT cell development in Id3-deficient mice

In the previous sections, we investigated the mechanisms that drive iNKT cell development in Id2/Id3-double deficient mice. Interestingly, Id3-deficient mice have been described to support the expansion of both γδNKT and iNKT cells.^[19, 47]^ We have also previously shown that varying activity levels of E proteins can differentially influence lineage outcomes between γδNKT and iNKT cells.^[15]^ Additionally, recent publications have demonstrated the sharing of transcriptional programs between these innate-like lineages, particularly between γδNKT and NKT2 cells.^[48, 49]^ Given the transcriptional and functional similarities between these two innate-like populations, and their expansion in Id-deficient mice, we wanted to examine if Id proteins suppressed these lineages equally or differentially, particularly in the developmental window prior to TCR selection. We therefore tested lineage competition in Id3-deficient mice by eliminating y5 lineage development and expansion.

We found a significant increase in the iNKT population in Id3^−/−^TCRδ^−/−^ mice as compared to Id3^−/−^ mice (Figure 2a-c). There was a modest but significant increase in iNKT lineage-committed stage 0 cells in Id3^−/−^TCRδ^−/−^ mice as compared to Id3^−/−^ mice (Figure 2d, e). It has been reported that iNKT cells and γδNKT cells compete for a thymic niche, based on the reduction in iNKT cells upon expansion of γδNKT cells.^[50]^ In contrast, another study has reported that a reduction in iNKT cells does not lead to a corresponding increase in γδNKT cells.^[51]^ In order to separate ongoing T cell development from homeostatic expansion associated with void space, we examined pre-weaning pups that have not yet undergone full expansion and stabilization of the thymic architecture. A large increase in the iNKT population was observed again in pre-weaning age Id3^−/−^TCRδ^−/−^ mice (Figure 2f, g). Age matched TCRδ^−/−^ mice, which lack total γδ T cells but are sufficient in Id3, also did not show any corresponding increase in iNKT cells. Overall, this data suggests that Id proteins control the outcome of the lineage competition between γδNKT and iNKT cells. Since the former arise at the DN3 stage and the latter at the DP stage, it also indicates that an early lineage checkpoint may be shared between them.

**Figure 2.**
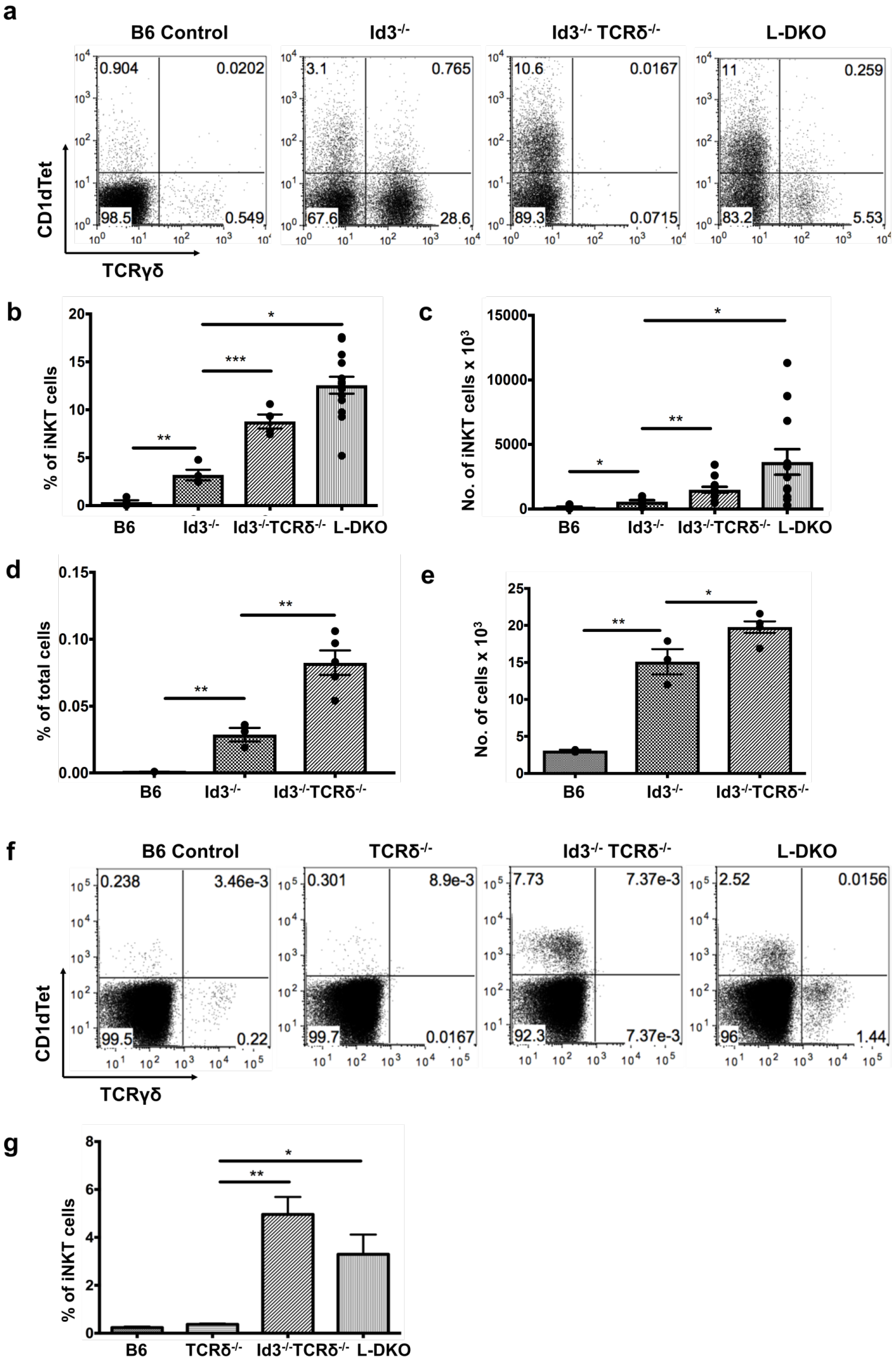
Absence of γδ T cells amplifies iNKT cell population in Id3^−/−^ mice. (a) Representative distribution of iNKT (CD1dTet+) versus γδ T cells in 20 day old B6 (n=4), Id3^−/−^ (n=4), Id3^−/−^TCRδ^−/−^ (n=4) and L-DKO (n=4) mice. (b) Percentage and (c) numbers of iNKT cells in 3-5 weeks old B6 (n=5), Id3^−/−^ (n= 7), Id3^−/−^TCRδ^−/−^ (n=13) and L-DKO (n=14) mice. (d) Percentage and (e) numbers of stage 0 iNKT cells in 2-5 weeks old B6 (n=3), Id3^−/−^ (n=3) and Id3^−/−^TCRδ^−/−^ (n=5) mice. (f) Representative distribution of iNKT (CD1dTet^+^) versus γδ T cells in 13 day old B6 (n=3), TCRδ^−/−^ (n=3), Id3^−/−^TCRδ^−/−^ (n=5) and L-DKO (n=4) mice. (g) Percentage of iNKT cells in 13 day old mice shown in (f). Error bars represent SEM. Statistical significance is represented by p values (* < 0.05, ** < 0.005, *** < 0.0005, n.s > 0.05).

### E2A drives a gene network that promotes iNKT and innate-like fate in the absence of Id proteins

Since both iNKT and γδNKT cell populations were amplified in the absence of Id protein(s), and we identified E2A as a potential regulator of iNKT-specific transcription factors, we wanted to determine whether E2A controls the transcription programs critical for either or both of these innate-like T cell lineages. We have previously characterized the predominantly enhanced γδNKT development in Id3^−/−^ mice, and iNKT development in L-DKO mice.^[15]^ Therefore, we performed RNA-Seq analysis of Id3^−/−^ γδ (predominantly γδNKT cells), L-DKO iNKT and L-DKO DP cells, and their wild type (WT) counterparts. Interestingly, both Principal Component Analysis (PCA) and clustering analysis highlighted transcriptional similarity between L-DKO iNKT and Id3^−/−^ γδNKT cells (Figure 3a, b). There were 1611 and 1865 genes dysregulated by at least two fold in L-DKO iNKT and Id3^−/−^ γδNKT cells respectively, as compared to their WT controls (Figure 3c). Among these, more than 400 genes shared similar expression patterns in the two cell types (Figure 3c, d). Pathway analysis of dysregulated genes revealed involvement of several pathways, including IL2/STAT5, p53 and MTORC signaling (Figure 3e). We also found enrichment of genes that were dysregulated in T-to-natural killer (ITNK) cells upon Bcl11b deletion (Figure 3e)^[52]^.

**Figure 3.**
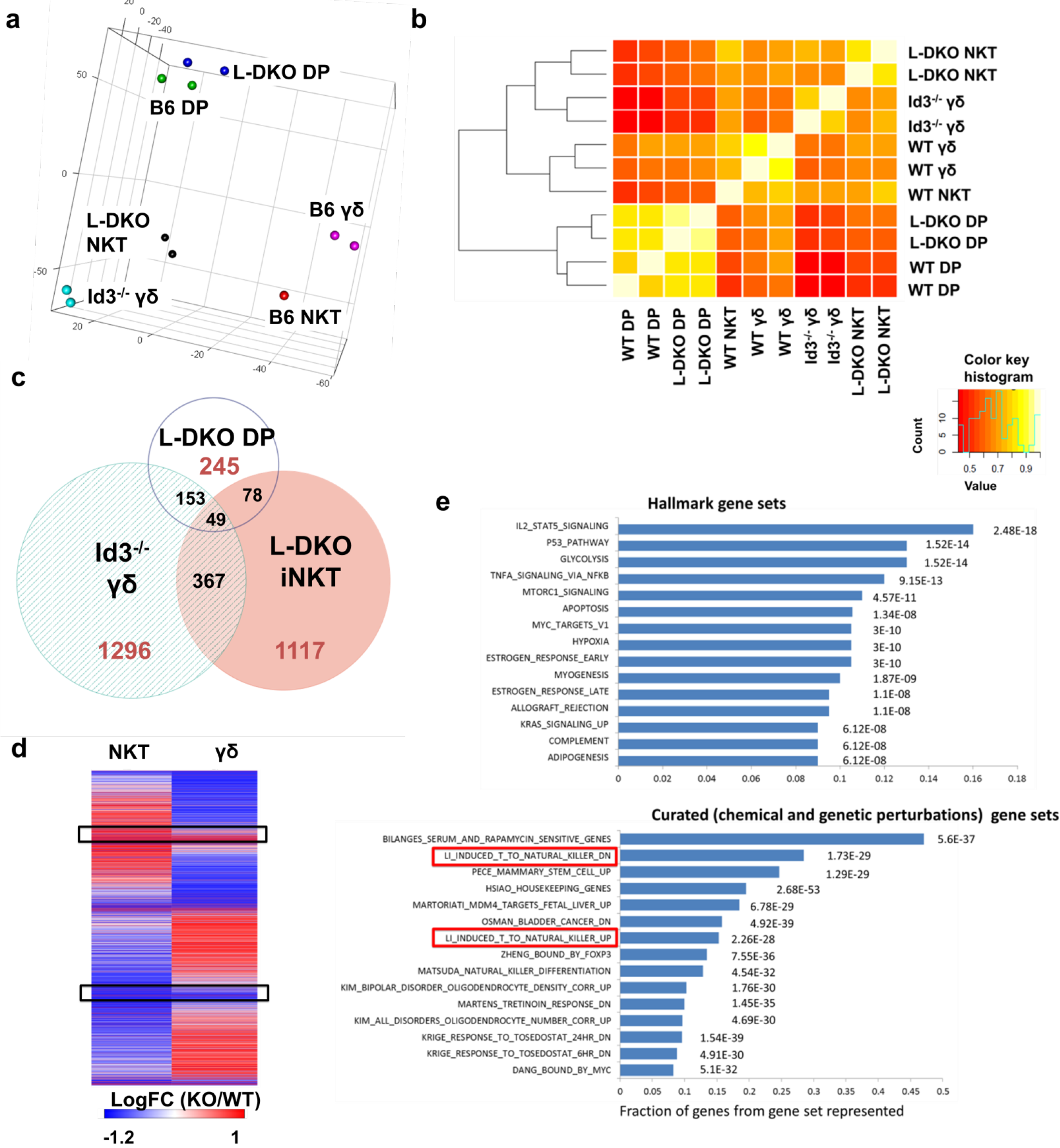
Transcriptional dysregulation in iNKT and γδNKT cells in the absence of Id proteins. (a) Principal Component Analysis of RNA-Seq expression data. (b) Hierarchical clustering analysis based on RNA-Seq expression patterns. (c) Genes found to be differentially expressed (fold change > 2) in L-DKO DP, L-DKO iNKT and Id3^−/−^ γδ samples, as compared to their WT counterparts. Numbers indicate unique or shared gene dysregulation between samples. (d) Expression patterns of differentially expressed genes identified in (c) in L-DKO iNKT and Id3^−/−^ γδ T cells compared to WT iNKT and WT γδ T cells respectively. Genes with upregulation or downregulation in both cell types are highlighted. (e) Pathway analysis of genes that were more than 2 fold dysregulated in L-DKO iNKT or Id3−/− γδ T cells. Top pathways from hallmark and curated gene sets are displayed with the percentage of genes that were shared with the respective pathways, along with the p values. Gene sets representing upregulation or downregulation of genes in T-to-natural killer (ITNK) cells upon Bcl11b deletion, as curated by Msigdb (gene sets M2088 and M2086), are highlighted in red.

In order to specifically identify any dysregulation of innate-specific genes, we decided to filter differentially expressed genes in L-DKO iNKT and/or Id3^−/−^ γδNKT cells, i.e. a total of 3060 genes, against an innate-like gene signature. We first searched for the reference innate-like signature genes from publicly available Immgen data.^[53]^ We hypothesized that the genes that are upregulated or downregulated significantly in both iNKT and γδNKT cells over other T cell populations in the thymus, would be unique to these innate-like lineages and most likely be important for their development and/or function. Upon TCR selection, the divergence of innate-like lineages from conventional T cells is primarily represented by CD44^−^NK1.1^−^ stage 0/1 iNKT cells and CD24^hi^ immature γδNKT cells. These cells are also phenotypically most similar to the expanding populations in Id-deficient mice. Therefore, we compared the gene expression in these WT innate-like T cells against multiple WT conventional T cell populations, including DN3a, DN3b, DN4, DP, post-selection CD4SP and thymic γδ T cells. This analysis resulted in 189 signature genes which are significantly overexpressed or repressed in both WT iNKT and γδNKT populations. We also added 7 other genes to this list that are reported in literature to be important for their development, but are not significantly overexpressed or repressed in these populations (**Supplementary Table I**). When this final innate-like gene set was compared to the genes identified in our RNA-Seq analysis, we found more than 50% (111 genes) to be dysregulated by at least two fold in either or both of the cell populations in Id-deficient mice (Figure 4a, **Supplementary Table II**). Importantly, E2A directly bound to many (83 of 111 genes) of these differentially expressed signature genes. This data suggested that E2A plays a profound role in regulating the expression of genes critical for innate-like T cell development.

**Figure 4.**
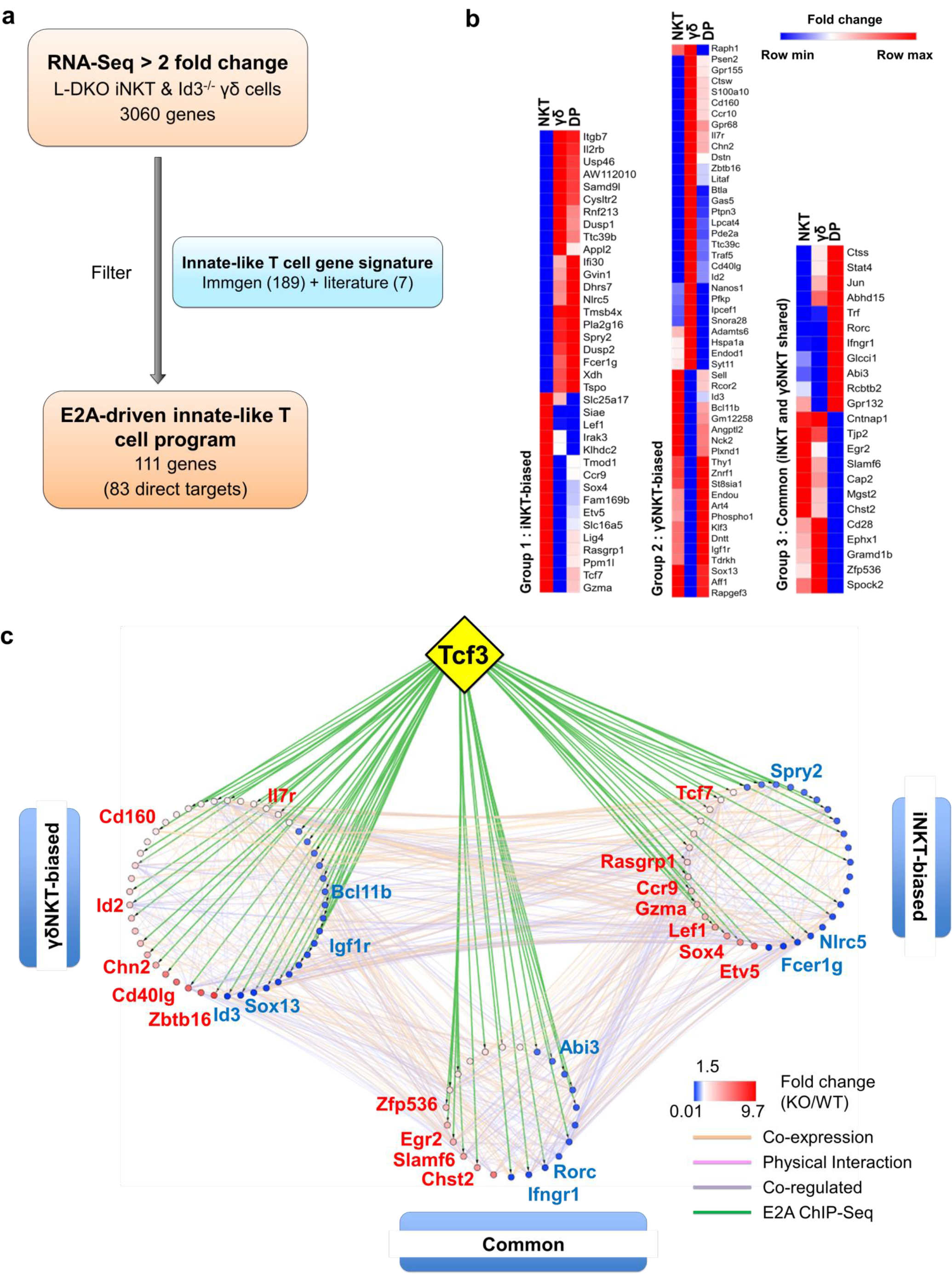
E2A regulates innate-like transcriptional programs in the absence of Id proteins. (a) Flowchart representing the analysis pipeline used to derive the E2A-driven innate-like T cell program depicted in (c). (b) Expression patterns (fold change in mutant cells compared to WT) of genes in the 3 groups shown in (c), based on more than 2 fold dysregulation in one or more cell types, and part of the innate-gene signature derived from WT Immgen data. (c) Network analysis combining RNA-Seq expression and E2A ChIP-Seq data for innate-like signature genes. Genes in the “γδNKT-“ or “iNKT-biased” groups are colored according to their expression in Id3^−/−^ γδ T cells and L-DKO iNKT cells respectively. Genes in the “common” group are represented by their average expression in d3^−/−^ γδ T cells and L-DKO iNKT cells. ChIP-Seq binding of E2A (encoded by Tcf3 gene) to gene targets is represented by green lines. Other interactions between gene targets, classified by GeneMania as co-expression, physical interaction or co-regulation, are represented by orange, pink and purple lines respectively.

In order to further delineate the role of E proteins in regulating the developmental programs of iNKT and γδNKT cells through these downstream mediators, we divided the 111 genes into 3 groups based on their expression profiles in L-DKO iNKT and Id3^−/−^ γδNKT cells (Figure 4b, **Supplementary Table II**). Groups 1 and 2 included “biased” genes that were upregulated or downregulated by a significantly larger magnitude in one population as compared to the other. On the other hand, group 3 included “common” genes that were significantly and similarly upregulated or downregulated in both innate-like populations. These categories broadly represent genetic programs that are potentially important for the lineage-specific development of iNKT (group 1) and γδNKT cells (group 2), as well as for overall innate-like T cell specification (group 3). By combining known interactions between these genes with our RNA-Seq and ChIP-Seq data, we created a network map with the three groups of genes demarcated (Figure 4c). The distribution of E2A targets across all three groups strongly reaffirmed the role of E2A in orchestrating innate-like T cell developmental programs. Our previous observation of diminished iNKT and γδNKT populations in Id2^f/f^Id3^f/f^E2A^f/f^HEB^f/f^ LckCre^+^ (or Q-KO) mice that lack E protein activity further supports the pivotal role of E proteins in the development of these cells.^[15]^

### Lack of Id proteins promotes innate-like T cell development independent of pre-TCR signaling

All observations thus far suggested the promotion of innate-like T cell fate by uninhibited E protein activity. The E2A occupancy of innate-like genes at DP cells and the lineage competition between iNKT and γδNKT cells further prompted us to consider the possibility that Id proteins may regulate the fate choice for innate-like T cells early in T cell development. We therefore tested the requirement of pre-TCR signaling in the development of innate-like T cells in Id-deficient mice. A functional pre-TCR signal at the DN3 stage is critical for conventional αβ T cell development. Mice deficient in pre-Tα, are therefore known to have restricted T cell development, with the majority of cells blocked at the DN stage.^[27]^ pTα^−/−^ mice are also known to completely lack iNKT cells.^[54]^ We crossed pTα^−/−^ mice to L-DKO to generate Id2^f/f^Id3^f/f^LckCre^+^ pTα^−/−^ (L-DKO pTα^−/−^) mice.

Despite the complete absence of iNKT cells in pTα^−/−^ mice, to our surprise, we found a robust iNKT population in L-DKO pTα^−/−^ mice (Figure 5a-c). These iNKT cells also expressed high levels of PLZF, which is closely associated with the innate-like lineage gene expression program (Figure 5d). It is known that pTα^−/−^ mice have an increase in γδ T cells,^[27]^ and L-DKO pTα^−/−^ mice showed a similar increase in the γδ population compared to WT mice (Figure 5e). However, the γδ T cells in L-DKO pTα^−/−^ mice were predominantly Vγ1.1^+^Vδ6.3^+^ and uniformly upregulated PLZF, reflecting a specific increase in innate-like γδNKT cells in these mice (Figure 5f-i).

**Figure 5.**
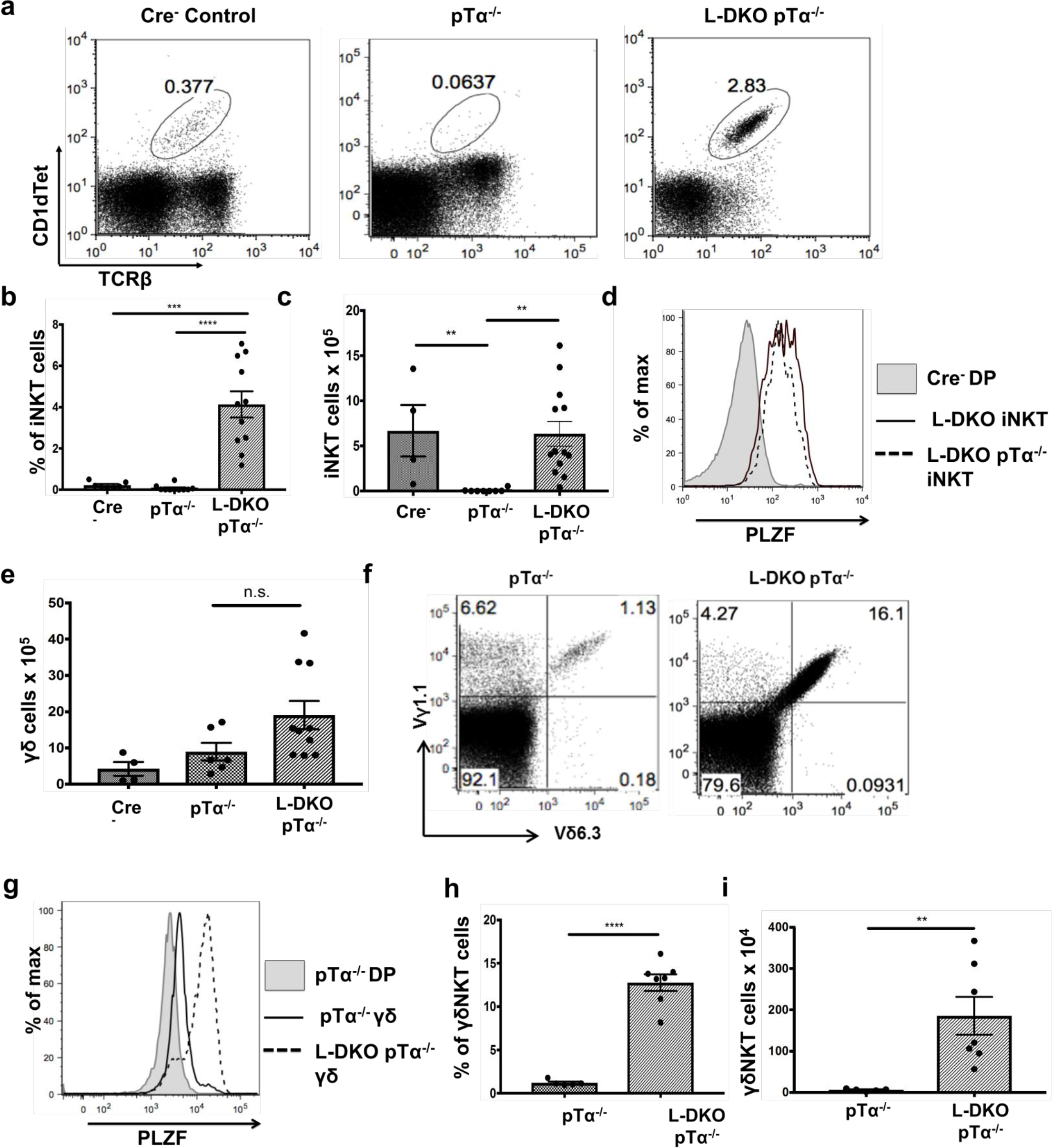
Lack of Id proteins promotes iNKT cell and γδNKT development independent of pre-TCR signaling. (a) Representative iNKT cells in 3-5 week old Cre^−^ (n=4), pTα^−/−^ (n=10) and L-DKO pTα^−/−^ (n=11) mice. (b) Percentage and (c) numbers of iNKT cells in 3-5 week old mice. (d) Representative PLZF expression levels in iNKT cells from 3-5 week old Cre-(n=3), L-DKO (n=3) and L-DKO pTα^−/−^ (n=4) mice. (e) Total number of γδ T cells in 3-5 week old Cre^−^ (n=4), pTα^−/−^ (n=6), L-DKO pTα^−/−^ (n=10) mice. (f) Representative γδNKT (Vγ1.1+Vδ6.3+) populations in 3-5 week old pTα^−/−^ (n=5), L-DKO pTα^−/−^ (n=7) mice. (g) Representative PLZF expression levels in γδ T cells from 3-5 week old Cre-(n=3), L-DKO (n=3) and L-DKO pTα^−/−^ (n=4) mice. (h) Percentages and (i) numbers of γδNKT cells in mice shown in (f).

As expected, L-DKO pTα^−/−^ mice still had a profound block in conventional T cell development due to the lack of pre-TCR signaling (Figure 6a, b). Interestingly, despite the pre-TCR block in L-DKO pTα^−/−^ mice, the deletion of Id proteins seemed to partially rescue the development of DP cells (Figure 6a, c). Upon careful investigation, however, we found that these DP cells upregulate PLZF (Figure 6d). These PLZF^hi^ DP cells, however, did not recognize the CD1d tetramer, indicating that they are not iNKT cells that aberrantly upregulate CD4 and CD8. Total thymocytes from L-DKO pTα^−/−^ mice also displayed a prevalent innate-like phenotype, as indicated by their PLZF expression pattern (Figure 6e). Thus, blocking conventional αβ T lineage development with pTα-deficiency revealed a pre-TCR independent pathway that drives innate-like T cell, including iNKT, lineage development in Id2/Id3-deficient mice.

**Figure 6.**
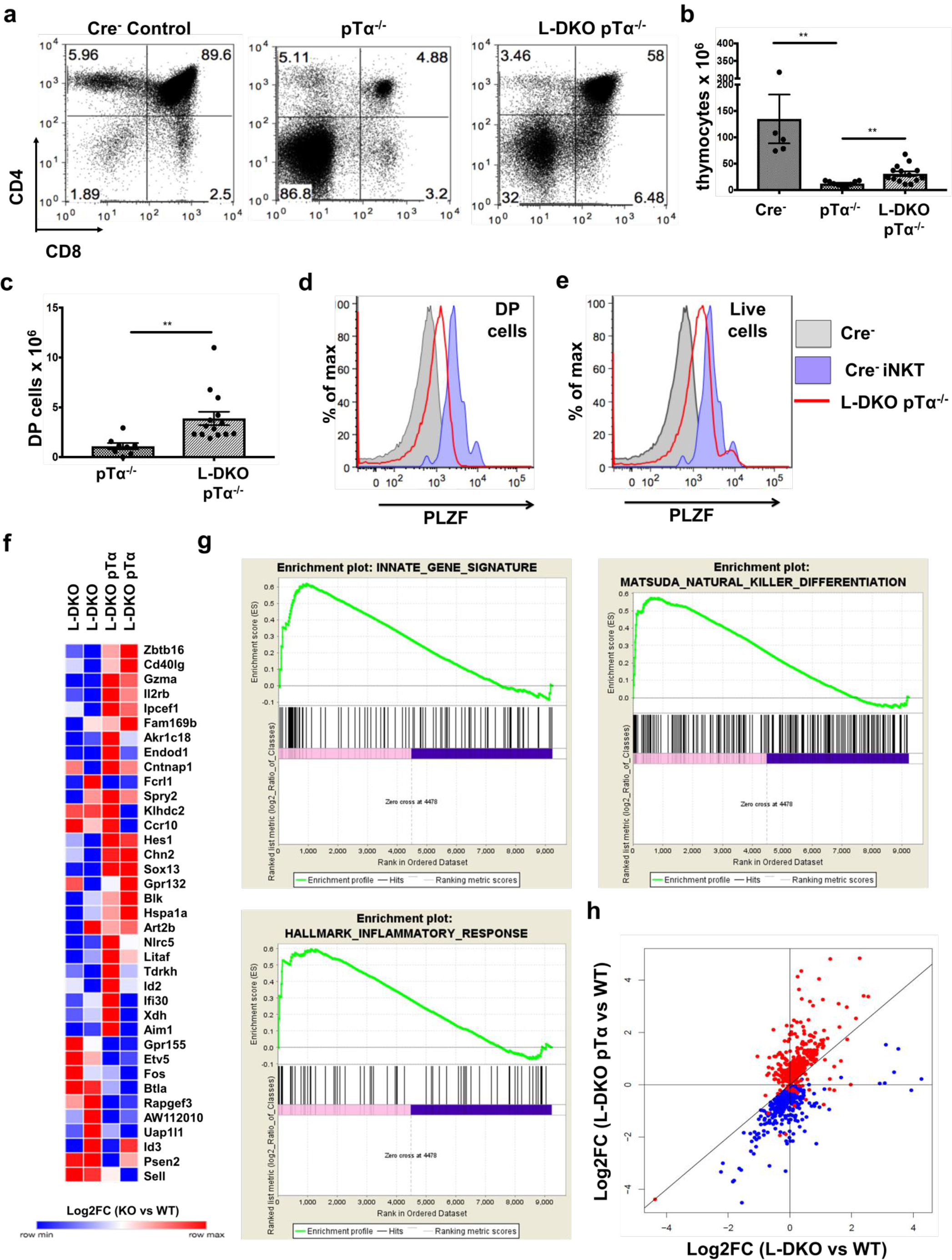
Adoption of innate-like transcription signature by DP cells in the absence of pre-TCR signaling and conventional T cells. (a) Representative thymocytes in 3-5 week old Cre^−^, pTα^−/−^ and L-DKO pTα^−/−^ mice as shown by CD4 and CD8 staining. (b) Total number of thymocytes in in 3-5 week old mice. (c) Number of DP cells in mice shown in (a). 5 to 13 mice included for each genotype shown in (a-c). (d, e) Representative PLZF expression levels in (d) DP cells and (e) total live cells from 3-5 week old Cre^−^ (n=3) and L-DKO pTα^−/−^ (n=4) mice. (f) Gene expression patterns (log2 fold change with respect to WT DP cells) for innate-like signature genes that are dysregulated by at least 2 fold in L-DKO or L-DKO pTα^−/−^ DP cells compared to WT DP cells. (g) Enriched gene sets in L-DKO pTα^−/−^ DP cells over L-DKO DP cells pertaining to innate-like gene signature (top left), iNKT development and maturation (top right) and inflammatory responses (bottom), using Gene Set Enrichment Analysis (GSEA). (h) Gene expression patterns (log2 fold change in L-DKO or L-DKO pTα^−/−^ DP cells with respect to WT DP cells) for genes that are positively (red) or negatively (blue) correlated with *Zbtb16* expression. Numbers indicate number of positively or negatively correlated genes in each quadrant or section. Error bars represent SEM. Statistical significance is represented by p values (* < 0.05, ** < 0.005, *** < 0.0005, n.s > 0.05).

### Initiation of an innate-like T cell transcriptional program in the absence of Id proteins and conventional T cell development

Our previous observation suggested that pre-TCR signaling and Id protein activity is necessary to enforce conventional T cell fate, such that the absence of both gave rise to predominantly innate-like T cell populations in the thymus. We further verified the innate-like phenotype of PLZF^hi^ DP cells in L-DKO pTα^−/−^ mice by RNA-Seq analysis. 28 genes out of the 196 genes in the innate-like gene signature defined previously (**Supplementary Table II**) were found to be upregulated by more than 2 fold in L-DKO pTα^−/−^ DP cells, as compared to WT DP cells (Figure 6f). The gene expression pattern for L-DKO DP was significantly different from L-DKO pTα^−/−^DP cells, including lower expression of key innate-like signature genes, such as *Zbtb16* and *Gzma*. Gene Set Enrichment Analysis (GSEA) verified enrichment of the innate-like gene signature, as well as genes associated with iNKT cell development and inflammatory responses in L-DKO pTα^−/−^ DP cells as compared to L-DKO DP cells (Figure 6g). RNA-Seq analysis of the PLZF^hi^ DP cells also revealed these cells had undergone TCRα rearrangement with a fairly broad V-J usage (data not shown).

Since PLZF is currently one of the only well-defined innate-like transcription factors, we decided to alternatively explore the innate-like transcription program by identifying genes that are positively or negatively correlated with PLZF expression. The idea is that genes with similar expression patterns can be expected to function together and/or be involved in similar biological processes. We computed pair-wise correlations between the expression of Zbtb16 and all other genes across WT, pTα^−/−^, L-DKO and L-DKO pTα^−/−^ DP cells (**Supplementary Table III**). A scatter plot of 465 positive and 460 negatively correlated genes revealed clear differences in expression patterns between L-DKO pTα^−/−^ and L-DKO DP cells (Figure 6h). Most genes upregulated in L-DKO pTα^−/−^ DP cells were positively correlated with Zbtb16, whereas most genes that were downregulated in these cells had negative correlation with Zbtb16. Interestingly, a significant fraction of positively correlated genes (308 genes) had a higher expression in L-DKO pTα^−/−^ DP cells. On the other hand, a significant fraction of negatively correlated genes (283 genes) had a higher expression in L-DKO DP cells. These analyses demonstrate the early initiation and adoption of an innate-like transcriptional program specifically in DP cells in L-DKO pTα^−/−^ mice. Therefore, we observe a complete diversion of cells towards innate-like T cell fate as a result of uninhibited E protein activity and lack of conventional T cells in the thymus. Overall, these results suggest that Id proteins are potent suppressors of innate-like T cell fate at the pre-TCR checkpoint.

## Discussion

In order to better understand the opposing roles of Id proteins in regulating conventional and innate-like T cell development, in this paper, we examined their role in suppressing innate-like T cells, including iNKT and γδNKT cells, prior to TCR selection. Our genetic studies clearly revealed a developmental behavior of the iNKT lineage that is inconceivable with our current understanding of these cells, whose lineage fate choice normally does not start until the TCR selection stage. Our finding represents the first case where iNKT lineage development can be distinguished from the conventional αβT lineage as early as the pre-TCR checkpoint, albeit under a unique genetic background. The combined genome-wide E2A binding and transcriptional data provides a direct explanation to the several previous reports of expansion of these innate-like populations in the absence of Id proteins. As more genes are increasingly reported to be important for innate T cell development and effector functions, our data also serves as a repository to examine the role of E2A in their regulation distinctly or commonly in both iNKT and γδNKT populations, particularly during early stages of their development.

Our E2A ChIP-Seq analysis of *Id2/Id3*-deficient (L-DKO) DP and iNKT cells placed E2A as either collaborator or upstream regulator of genes and transcription factors known to be important for iNKT cell development. The regulation of these gene targets at the DP stage, prior to the TCR-mediated selection of iNKT cells, indicated early lineage specification events for iNKT cells, as regulated by E2A. We also observed shared binding patterns of E2A in L-DKO DP and iNKT cells, with those previously documented in DN3 and DN4 cells with no Id deficiency.^[11]^ This suggests that several E2A targets identified in *Id*-deficient mice may also be similarly regulated in WT mice to promote innate-like T cell development.

It has been shown that the effector programs of iNKT2 cells closely resemble that of γδNKT cells,^[49]^ even though these two lineages are considered to independently diverge at the DP and DN3 stages respectively. The effector iNKT cells expanded in *Id2/Id3*-deficient mice most closely resembles the iNKT2 lineage.^[21, 46]^ Clustering analysis of our RNA-Seq data indicated transcriptional similarity between both WT and *Id*-deficient iNKT and γδNKT cells. The lineage competition between these innate-like populations revealed by Id3^−/−^TCRδ^−/−^ mice further hints towards the developmental lineage proximity between these two lineages. Due to the typically neonatally restricted development of γδNKT cells, it is unfeasible to verify this phenotype using bone-marrow chimeras.^[19]^ However, the increase in stage 0 iNKT cells in the absence of γδNKT cells suggests that these lineages might have shared innate-like precursors, whose lineage choice is regulated by Id proteins.^[55]^ This data also challenges the stochastic TCR-driven iNKT developmental model at the DP stage, and warrants re-examination of the lineage origin of iNKT cells.

Our study also revealed E2A-mediated transcription programs that support the development of γδNKT and iNKT lineages. This analysis identified E2A as an upstream regulator of genes critical for iNKT lineage differentiation, including *Zbtb16*, *Slamf6* and *Egr2* (Figure 3e).^[3, 4, 6, 23, 24, 56–59]^ Among other target genes, we also found a dramatic downregulation of *Bclllb* in γδNKT cells, which is needed to suppress a Natural Killer (NK) program in conventional T cells.^[60]^ This indicated a possible role for *Bclllb* in the lineage choice between conventional *γδ*T and γδNKT cell development. Genes that are associated with NTK1 and NKT17 cytokine profiles, such as *Ifngr1* and *Rorc*, were found to be significantly downregulated in both Id3” “ γδNKT and L-DKO iNKT cells. These findings demonstrate the involvement of E2A in preferentially driving γδNKT and NKT2 lineage development in the absence of Id proteins.

In this study, we also uncovered a pre-TCR independent pathway for the development of iNKT and innate-like T cells using our L-DKO pTα^−/−^ mice. Interestingly, the expanded innate-like phenotype in the L-DKO pTα^−/−^ mice was not limited to only iNKT and γδNKT cells, but also a novel innate-like PLZF^hi^ DP population. The early induction of an innate-like transcriptional program among all developing T cells in these mice with high E protein activity demonstrates the necessity of Id proteins in suppressing innate-like fate choice. The expanded innate-like populations in our *Id*-deficient mice allowed us to uncover the regulation of innate-like T cell lineage development prior to TCR selection. However, the divergence of iNKT and innate-like T cells from conventional T cells prior to TCR selection has also been proposed in other mouse models with physiological levels of E protein activity.^[55, 61]^ It is likely that the depletion of Id proteins unleashes the “early”, pre-TCR-independent developmental program for iNKT and other innate-like T cells, that can usually be observed in much smaller frequencies in other studies. Consequently, we have also observed heterogeneous innate-like αβ T cell lymphomas in Id2/Id3 deficient mice that are derived from iNKT, CD1dTet^−^ or T_FH_ cells.^[16, 62]^

PLZF undoubtedly plays key roles in innate-like T cell development and effector function. However, the transcriptional regulation of PLZF itself remains ambiguous. While some studies have found strong agonistic signaling to be sufficient to induce PLZF expression, ^[5, 24, 63]^ others have reported early and stable suppression of PLZF in conventional T cells, which can’t be reversed by TCR signaling alone.^[64]^ The function of PLZF itself, in promoting innate-like effector genes, has also been demonstrated to be independent of agonistic TCR signals.^[65, 66]^ This implies that unveiling the transcriptional regulation of PLZF and lineage specification of innate-like fate in developing T cells is critical for advancing our understanding of innate-like T cells. Our data suggests that Id-mediated inhibition of E2A is able to suppress the downstream activation of PLZF and innate-like transcription programs in majority of developing T cells as early as the pre-TCR selection step. This supports a layered, rather than a parallel, developmental structure that coordinates the distinct developmental fates of innate and conventional T cells in the thymus^[45]^.

Although innate-like T cells represent only a small fraction of the thymic population, their indispensable roles in mounting rapid immune responses in different contexts warrants a holistic understanding of the regulation of their concurrent development with conventional T cells in the thymus. In this study, we characterized E2A-driven transcriptional programs that promote innate-like T cell development prior to TCR selection and independent of pre-TCR signaling, which are otherwise suppressed by Id proteins. Not surprisingly, phylogenetic analysis of innate-like T cells and their associated transcription factors indicates that these cells emerged much earlier than conventional T cells in the course of evolution.^[45, 67]^ Hence, we propose that innate-like lineage specification precedes conventional αβ T cells in the thymus, and that evolutionary pressures necessitated *Id*-mediated suppression to ensure the predominance of conventional αβ T cells. Our data also suggests that the collaborative action of Id proteins and pre-TCR signaling delays the fate choice of iNKT and innate-like T cells to drive conventional T cell development in the thymus.

## Acknowledgments

We thank Drs. A. Lasorella and A. Lavarone (Columbia University) for sharing the Id2f/f strain. We thank the Duke Cancer Center Flow Cytometry Facility for assistance in cell sorting, Duke Cancer Center Sequencing Facility for assistance in Ion Torrent sequencing, UC San Diego IGM Genomics Centre for assistance in ChIP-Seq sequencing, Duke Center for Genomic and Computational Biology for assistance in RNA-Seq sequencing, and the NIH tetramer facility for providing the CD1d tetramer. We thank Drs. M. Krangel, M. Ciofani, B. Mathey-Prevot and Mr. Zachary Carico (Duke University) for their expertise and feedback.

## Footnotes

1. This work has been supported by National Institute of Health grants GM R01 GM059638, and 1 P01 AI102853 awarded to Dr. Yuan Zhuang.
2. Id : Inhibitor of DNA binding
3. iNKT - invariant Natural Killer T cell
4. CD 1dTet - CD 1 d tetramer
5. L-DKO - Id2^f/f^Id3^f/f^LckCre^+^
6. TKO - Id2^f/f^Id3^f/f^LckCre^+^CD 1d^−/−^
7. PLZF - Promyelocytic Leukemia Zinc Finger
8. WT - Wild Type

